# Optogenetic and transcriptomic interrogation of enhanced muscle function in the paralyzed mouse whisker pad

**DOI:** 10.1101/473298

**Authors:** Thomas J. Vajtay, Akhil Bandi, Aman Upadhyay, Mavis. R. Swerdel, Ronald P. Hart, Christian R. Lee, David J. Margolis

## Abstract

The functional state of denervated muscle is a critical factor in the ability to restore movement after injury- or disease-related paralysis. Here we used peripheral optogenetic stimulation and transcriptome profiling in the mouse whisker system to investigate the time course of changes in neuromuscular function following complete unilateral facial nerve transection. While most skeletal muscles rapidly lose functionality after lower motor neuron denervation, optogenetic muscle stimulation of the paralyzed whisker pad revealed sustained increases in the sensitivity, velocity, and amplitude of whisker movements, and reduced fatigability, starting 48 h after denervation. RNA-seq analysis showed distinct regulation of multiple gene families in denervated whisker pad muscles compared to the atrophy-prone soleus, including prominent changes in ion channels and contractile fibers. Together, our results define the unique functional and transcriptomic landscape of denervated facial muscles, and have general implications for restoring movement after neuromuscular injury or disease.

**New & Noteworthy:** Optogenetic activation of muscle can be used to non-invasively induce movements and probe muscle function. We used this technique in mice to investigate changes in whisker movements following facial nerve transection. We found unexpectedly enhanced functional properties of whisker pad muscle following denervation, accompanied by unique transcriptomic changes. Our findings highlight the utility of the mouse whisker pad for investigating the restoration of movement after paralysis.

## INTRODUCTION

The functional state of denervated muscle is critical for the restoration of movement after paralysis caused by chronic injury or disease. Functional changes in muscle are well known after lower motor neuron denervation for many types of muscle (Thesleff 1974, Wu et al 2014), with loss of function, atrophy and reduced capacity for movement being the most common. Gain-of-function changes can also occur in parallel, including increased excitability and hypersensitivity to the neurotransmitter acetylcholine (Jones & Vrbova 1974, Lomo & Rosenthal 1972, Sellin & Thesleff 1980). However, gain-of-function changes do not counteract the atrophy and loss of function in most muscles, which presents major problems for generation of contractile force and the ability of muscle to respond to restorative interventions. Furthermore, available clinical strategies to stimulate neuromuscular function generally target a partially intact nerve instead of direct stimulation of the muscle because it is difficult to produce well-controlled, graded movements using electrical stimulation of muscle (Doucet et al 2012, Griffin & Kim 2011, Ho et al 2014, Peckham & Knutson 2005). Non-invasive, direct stimulation of paralyzed muscle that produces naturalistic movements is highly desirable but has not been achieved using traditional methods.

Our previous work established a method for non-invasive control of whisker movements in mice using transdermal photostimulation of the optogenetic actuator channelrhodopsin-2 (ChR2) expressed in whisker pad muscles (Park et al 2016). Spot illumination of the rostral whisker pad with blue light in Emx1-ChR2 mice, as used in the previous and current work, evokes whisker protractions by activating extrinsic protractor muscles pars media superior and pars media inferior of *M. nasolabialis profundus* and also intrinsic follicular muscles (Haidarliu et al 2015, Haidarliu et al 2010, Park et al 2016), collectively referred to here as “whisker pad muscles”. In the present study, we used ChAT-ChR2 and Emx1-ChR2 mice to evoke whisker movements via stimulation of the facial motor nerve (cranial nerve VII) or the whisker pad muscles, respectively. This allowed us to investigate the functional changes that occur in nerve and muscle after the paralysis of whisker movements caused by facial nerve transection.

One recent study used optogenetic muscle stimulation in the hindlimb triceps surae after sciatic nerve lesion to demonstrate dramatic atrophy and loss of function (Magown et al 2015), consistent with classic studies in this system (Nelson 1969), that could be attenuated by daily optogenetic activation. We considered it possible that whisker pad muscles undergo distinct denervation-induced changes compared to other muscle types, in part because whisker pad muscles are composed of distinctive fiber type (Haidarliu et al 2010, Jin et al 2004). While the rodent whisker system has been used for studies related to recovery of function and reinnervation (Hadlock et al 2005, Heaton et al 2014), the functional state of denervated whisker pad muscles and their capacity to support evoked movements has remained unclear, in part because non-invasive methods to probe muscle function have only recently been established.

We longitudinally tracked whisker movements evoked by optogenetic nerve or muscle stimulation up to 10 d after facial nerve transection. While nerve-evoked responses degraded by 1 d, muscle-evoked whisker protractions showed dramatic increases in sensitivity, amplitude, velocity, and reduced fatigability. Furthermore, RNA-seq analysis of denervated muscle at days 1, 3 and 7 revealed transcriptome-level changes in ion channels and contractile fibers (among others), with many striking differences compared to published data from the atrophy-prone soleus muscle. Our results could lead to development of treatment strategies for restoring function in various types of paralyzed muscle.

## METHODS

### Subjects and surgical preparation

Procedures were approved by Rutgers University Institutional Animal Care and Use Committee. Mice were purchased from Jackson Labs (Bar Harbor, ME) and bred in house. ChR2 expression was driven in nerve crossing Ai32 Cre-dependent ChR2 reporter mice (B6;129S- Gt(ROSA)26Sortm32(CAG-COP4*H134R/EYFP)Hze/J; Jax 012569) with ChAT-Cre mice (B6;129S6-Chattm2(cre)Lowl/J; Jax 006410) to produce “Chat-ChR2” mice. Chat-ChR2 mice express ChR2 in cholinergic neurons, including motoneurons (Rossi et al 2011). Ai32 mice were crossed with Emx1-Cre mice (B6.129S2-Emx1tm1(cre)Krj/J; Jax 005628) to produce “Emx1-ChR2” mice (Madisen et al 2012) that express ChR2 in whisker pad muscles (Park et al 2016). Additional groups of mice were used for pharmacological effects on fasciculations (n=5) and RNA-seq (n=9) experiments (below). Emx1-ChR2 mice on the day of lesion were postnatal day (P) 52 or P55. Additional Emx1-ChR2 mice used for ChR2 expression measures were P75 on the day of lesion. ChAT-ChR2 mice were P276 on the day of lesion. Mice were median age P224 (range P159-224) for RNA-seq experiments. Both male and female mice were used, with weights between 19 and 38 g.

For optogenetic stimulation, adult mice were implanted with a head post affixed to the skull with dental cement in a surgical procedure under isoflurane anesthesia, as described (Lee & Margolis 2016, Park et al 2016), and allowed to recovery for at least 5 d. After baseline measurements were taken, facial nerve transection was performed following (Olmstead et al 2015) in a second surgery. Briefly, mice were anesthetized with isoflurane (4% induction, 2% maintenance) and depilatory cream was applied to an approximately 1×1 cm area caudal to the whisker pad between the eye and jaw. A dorsoventral skin incision was made using spring scissors, and the buccal and superior marginal branches of the facial nerve exposed before transection with spring scissors. The wound margin was closed using cyanoacrylate glue and mice allowed to recover in the home cage. Unilateral whisker movement paralysis was visually verified.

### Stimuli and video recording

Light stimuli and recording methods followed our previous work (Park et al 2016). Briefly, a blue 460 nm high-power LED (Prizmatix) was coupled to a 200 µm diameter optical fiber (Thorlabs), producing a 2-3 mm spot of light on the face. Video of whiskers was recorded at 500 frames/s with a high-speed CMOS camera (PhotonFocus, DR1) under infrared illumination (Advanced Illumination). 8 mW stimuli of varying duration were triggered using TTL pulses sent by Arduino microcontroller to the LED current controller. Movies of 3 s duration were recorded for each trial with a 5 s or 15 s inter-trial interval for Emx1-ChR2 and ChAT-ChR2 mice respectively. Sequential trials increased in stimulus duration up to 1000 ms, at which point the sequence was repeated. There were 8 replicates of each stimulus duration for each mouse. To identify the position of the nerve in ChAT-ChR2 mice, the buccal branch of the facial nerve (cranial nerve VII) was targeted with the light spot at a position between the stylomastoid foramen, ventral to the ear canal and caudal to the whisker pad. Different illumination positions in this region were tested for each subject to optimize the evoked whisker protraction. To test effects on fasciculations, dantrolene (1 mM) was applied subcutaneously to the lesioned (right-side) whisker pad. The placement of LED for muscle stimulation in Emx1-ChR2 mice followed Park et al (2016) to target the rostral protraction or caudal retraction area.

### Whisker tracking and measurements

Frame-by-frame whisker angle was extracted using routines in Matlab and Python (Clack et al 2012). Video artifacts were corrected using ImageJ if the whiskers were mislabeled. Time series data from individual whiskers were further analyzed in Matlab, including maximum amplitude, velocity (peak of first derivative after Savitzky-Golay smoothing of single trials) and time-to-peak. Persistence index, a measure of fatigability, was calculated for 1000 ms stimuli as the ratio of response amplitude at 1000 ms to peak amplitude. Spectral analysis of fasciculations was performed using the fast Fourier transform (FFT) function within Matlab. Data from trials were averaged within and then between mice. Statistics were calculated using linear mixed models using the lme4 R package. Post hoc testing was done with paired contrasts via the multcomp R package and p values were adjusted for multiplicity with the Holm-Bonferroni method.

### RNA-seq

A separate group (n=9 mice) was used for tissue extraction for RNA-seq. Nerve transection was performed as above, with the addition of a sham control on the opposite whisker pad, where the skin was opened and re-closed but the nerve was not lesioned. At 1, 3, and 7 d post-lesion, mice were anesthetized with isoflurane (4% induction) and the fur trimmed to the level of the skin, then killed by cervical dislocation. Skin and muscle tissue was harvested bilaterally from lesioned and intact rostral whisker pads using a 3 mm circular biopsy punch at the site of the rostral protraction area used for optogenetic stimulation. Biopsied tissue was flash-frozen in liquid nitrogen and ground with ceramic mortar and pestle prior to Trizol extraction. Sequencing libraries were prepared with the Illumina TruSeq RNA Sample Preparation Kit v2 and then RNA-seq was performed with the Illumina NextSeq 500. qPCR was used to verify selected genes. Comparisons were made between lesioned and sham-control sides as well as a soleus dataset from GEO: GSE58669 (Macpherson et al 2015) after replicating the latter analysis in our framework to determine significant genes. RNA-seq data have been uploaded to NIH GEO with accession number GSE121590.

Illumina bcl2fastq2-v2.17.1.14 software was used for basecalling. Sequences were de-duplicated with ParDRe version 2.1.5-PC170509. Adapter sequences were trimmed and QC performed with Fastp version 0.12.2. De-duplicated, trimmed sequences were aligned with ENSEMBL GRCm38 cDNA transcripts using Kallisto version 0.43.1 with 100 bootstrap simulations (-b 100). Kallisto transcript counts were loaded into Sleuth version 0.30.0. The linear model for likelihood ratio test (LRT) was ∼ days:days + injury. A Wald test (WT) was evaluated for each day by separately loading in respective data and comparing sides. The significant WT transcripts were then filtered with the differentially expressed transcripts from the LRT and limited to only those transcripts with at least a 2 FC. Enriched GO terms were determined by creating a list of the gene ID’s of all 2 FC transcripts calculated from the WT, sorted by FC, and then analyzed with clusterProfiler for both cellular component and biological process enriched GO terms. Enriched GO terms for whisker 3 d were: GO:0014704; GO:0031674; GO:0043292; GO:1902495; GO:0005892; GO:0034702; GO:0030018; GO:0042641; GO:0000151; GO:0001518. Enriched GO terms for soleus 3 d were GO:1902495; GO:0005901; GO:0034702; GO:0008076; GO:0014704; GO:0042383; GO:0043292; GO:0043235; GO:0016528.

The transcripts from significant GO terms were then compared between whisker and soleus data in heatmaps. Normalized transcripts per million were used to calculate a z-score of the injured expression levels for each day separately and then only those transcripts that were differentially expressed in at least one of the data sets were isolated. To further isolate transcripts of interest, we then limited the selection to only those transcripts that had polar expression between whisker and soleus 3 d.

### Quantitative real-time PCR (qPCR)

qPCR was used to validate RNA-seq results and to test for changes in ChR2 expression Supplemental Fig. 4 (goo.gl/sb75Qd). Total cellular RNA was reverse transcribed into cDNA, then assayed by qPCR using SYBR Green PCR Master Mix (Life Technologies) following the manufacturer’s recommendations. Primers were designed to be specific using Primer-BLAST. Beta-Actin was used to normalize expression levels to obtain relative quantity (RQ) values. Forward and reverse primers are listed in Supplemental Fig. 4 (goo.gl/sb75Qd).

### Data availability

Data and code to recreate figures and statistics are available at https://github.com/margolislab/Vajtay_Bandi_2018. Source data are available upon reasonable request. RNA-seq dataset is available at https://www.ncbi.nlm.nih.gov/geo/query/acc.cgi?acc=GSE121590.

## RESULTS

We aimed to determine the functional and transcriptomic state of denervated (paralyzed) whisker pad muscle. We used longitudinal optogenetic methods to investigate the time course of whisker movements evoked by nerve or muscle stimulation up to 10 d after facial nerve transection, and in parallel, generated RNA-seq datasets on 1, 3 and 7 d.

### Rapid loss of nerve-evoked movements

We first performed optogenetic stimulation of the facial nerve in ChAT-ChR2 mice to determine the time course of functional denervation after nerve transection (**Fig. 1A**). Prior to nerve transection, transdermal illumination of the buccal branch caudal to the whisker pad evoked brief, high-velocity whisker protractions (peak amplitude, 8.77 ± 1.55°; peak velocity, 940 ± 161°/s; time to peak, 16.2 ± 0.93 ms; half width, 13.8 ± 1.16 ms SEM; n=6 mice). After facial nerve transection caudal to the illumination site, whisker movements could still be evoked at 0.5 and 6 h post-lesion, but were abolished by 24 h (**Fig. 1B**). Responses to brief (5 ms) or prolonged (1000 ms) 8 mW illumination were similar in both amplitude and velocity, suggesting that the facial nerve responds to optogenetic stimuli in an all-or-none fashion to suprathreshold stimuli. Analysis of group data from n=6 mice indicated that peak amplitude and peak velocity decreased in parallel over 24 h (**Fig. 1C, left, middle**). At the 6 h time point, peak amplitude decreased compared to baseline in 5/6 mice, and was fully abolished in 6/6 mice by 24 h (ANOVA of peak amplitude and peak velocity over days, F=31.16 and 37.88, p=0.0008 and 0.0005). The time-to-peak and half-width of the responses remained similar at the 6 h time point until responses were abolished at 24 h (**Fig. 1C, right**) (ANOVA of time-to-peak and half-width 1000 ms light response over days, F=58.19 and 22.07, p=1.7e-8 and 1.5e-6). During the course of nerve degradation (at 6 h), response duration families for peak amplitude and peak velocity showed similar overall shape and stimulus threshold (**Fig. 1D, left, middle**). Time-to-peak and half-width measures were constant when probed with stimuli of varying duration (**Fig. 1D, right**), indicative of stereotyped all-or-none nature of whisker movements evoked by suprathreshold optogenetic nerve stimulation. Together, these results indicate that the distal cut end of the facial nerve remains viable for at least 6 h, retaining its ability to evoke whisker movements in response to optogenetic stimulation, but fully degrades by the 24 h time point. Thus, whisker pad muscles are functionally denervated by 24 h after facial nerve transection.

**Figure 1.**
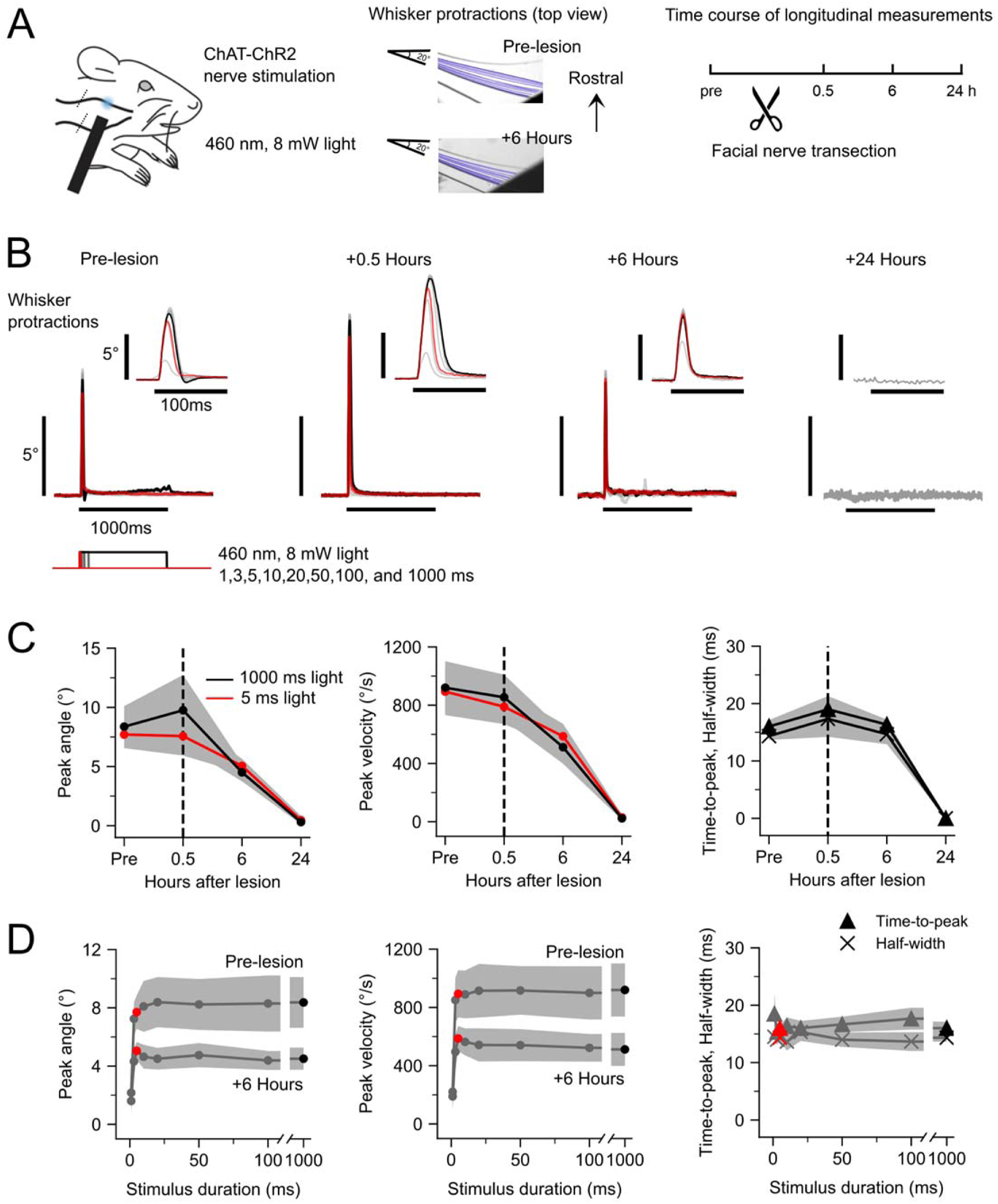
Functional degradation of facial nerve. **(A)** Optogenetic nerve paradigm. Left: Schematic of transdermal facial nerve illumination in ChAT-ChR2 mice. Position of optogenetic illumination indicated with blue spot. Position of nerve transection indicated with diagonal lines crossing buccal and marginal nerve branches caudal to illumination site. Middle: Frame-by-frame position of one tracked whisker in an example subject in response to a 1000 ms, 8 mW pulse of 460 nm LED illumination pre-lesion and 6 h post-lesion. Right: Experimental timeline. **(B)** Whisker protractions evoked by optogenetic nerve stimulation from an example mouse at various times relative to nerve transection. Each trace is the average of 10 responses to a different duration of illumination at 8 mW. Stimuli are 1, 3, 5, 10, 20, 50, 100, 1000 ms, with 5 ms (red) and 1000 ms (black) highlighted. Insets show the same traces on an expanded time scale. **(C)** Time course of peak amplitude (left), peak velocity (middle), time-to-peak and half-width (right) of whisker protractions in response to 5 ms (red) and 1000 ms (black) light stimuli. In all cases, responses were significantly abolished by 24 h post-transection (see text for further statistics). Values are mean ± SEM; n = 6 mice. **(D)** Pre-lesion and 6 h response duration families for peak amplitude (left) and peak velocity (middle). Pre-lesion response duration families for time-to-peak and half-width (right). Red circle, 5 ms; black circle, 1000 ms stimuli. Values are mean ± SEM; n = 6 mice.

### Spontaneous fasciculations

We observed spontaneous small-amplitude whisker twitches on the lesioned side after nerve transection (2.45 ± 0.54° peak-to-peak amplitude, n=9 mice), reflecting fasciculations (or fibrillations) of the denervated muscle fibers (Heaton & Kobler 2005, Salafsky et al 1968, Thesleff & Ward 1975, Wu et al 2014) (**Fig. 2A**). Fasciculation strength, as indicated by spectral analysis, increased in 7/9 mice at 2 d and 9/9 mice at 3 d post-lesion, and persisted through 10 d (**Fig. 2B**) (ANOVA of average power of fasciculations at 5-8 Hz over days, F=36.3, p<2.2e-16). In a separate group of mice, we tested the effects of the ryanodine receptor 1 (RyR1) antagonist dantrolene (1 mM) by subcutaneous whisker pad administration before and 7 d after nerve transection, and found that dantrolene abolished fasciculations (**Fig. 2C**) (ANOVA of average power of fasciculations at 5-8 Hz of DMSO, Dantrolene, and untreated mice before and 7 d after lesion, F=7.98, p=0.002; Paired contrasts of 7 d Dantrolene versus 7 d DMSO vehicle, t=2.95, p=0.04; n=5 mice), indicating a postsynaptic muscle origin. These results suggest that denervated whisker pad muscles undergo changes in intrinsic excitation-contraction coupling or excitability, raising the question of their capacity to sustain evoked movements.

**Figure 2.**
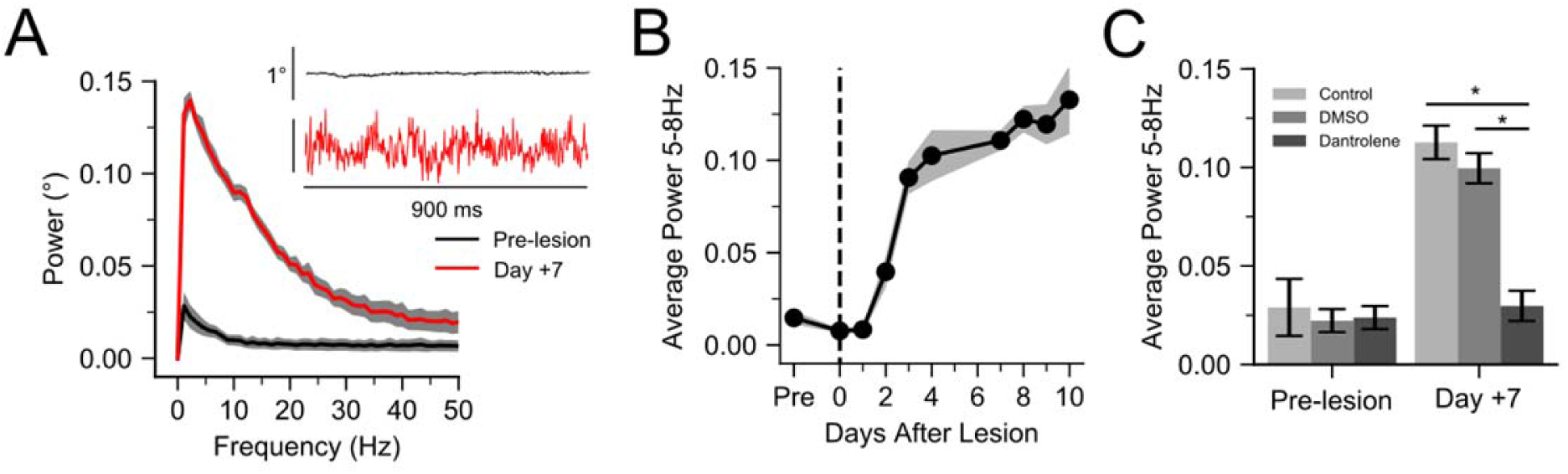
Muscle origin of fasciculations. **(A)** Power spectra of spontaneous whisker fasciculations at baseline and 7 d post-lesion. Inset, examples of spontaneous whisker movements at baseline and 7 d. **(B)** Time course of fasciculation strength (mean ± SEM 5-8 Hz power; n = 9, Emx1-ChR2 mice). **(C)** Effects of RyR1 antagonist dantrolene (1 mM). Bar graphs show mean ± SEM of 5-8 Hz power (n = 5, Emx1-ChR2 mice); * indicates p < 0.05.

### Denervation-induced enhancement of optogenetic muscle-evoked whisker movements

To determine functional properties of denervated whisker pad muscle, we used direct optogenetic muscle stimulation in Emx1-ChR2 mice following facial nerve transection (**Fig. 3A**). Emx1-ChR2 mice express ChR2 in whisker pad muscles (Park et al 2016) in addition to the well-known expression in excitatory cortical neurons (Madisen et al 2012). We performed longitudinal measurements of the same subjects, similar to the nerve experiments in figure 1, but over a longer time course up to 10 d. Transdermal illumination of the rostral protraction area (Park et al 2016) of the whisker pad under baseline conditions evoked robust whisker protractions (peak amplitude, 13.9 ± 1.6°; peak velocity, 414.8 ± 75.8°/s; 97.85 ± 14.26 ms time to peak; 368.17 ± 86.80 ms half width, n=9 mice) that adapted (or fatigued) during prolonged illumination. After facial nerve transection, peak amplitude and peak velocity of evoked movements began to increase on subsequent days, as shown for an example subject (**Fig. 3B**). Analysis of group data (n=9 mice) showed prominent increases in peak amplitude, peak velocity, and persistence index (a measure of reduced fatigability, see below) starting at 2 d post-transection and lasting throughout the 10 d time course (**Fig. 3C**) (ANOVA of peak amplitude and peak velocity over days, F=14.93 and 30.82, p=0.0001 and 0.005). For example, at 7 d, the peak amplitude of movements in response to 1000 ms illumination increased in 8/9 mice to on average 146% of baseline values (13.88° to 20.32°; ANOVA of 1000 ms peak amplitude responses over days, F=4.55, p=0.0001), and the responses to 5 ms stimuli increased by 505% of baseline (**Fig. 3C, left**) (2.75° to 13.90°; ANOVA of 5 ms peak amplitude responses, F=45.01, p<2.2e-16). Peak velocity of evoked whisker movements showed similarly dramatic increases. For example, at 7 d post-lesion, the peak velocity of movements in response to 5 ms stimuli increased until it was indistinguishable from the velocity of movements evoked by 1000 ms stimuli (**Fig. 3C, middle**) (834.17°/s for 5 ms stimuli, 991.49°/s for 1000 ms stimuli), indicating increased sensitivity of denervated whisker pad muscle. We further analyzed the fatigability (or adaptation) of evoked movements by measuring the persistence index of responses to 1000 ms stimuli (the ratio of final amplitude to peak amplitude; see inset in **Fig. 3C, right**). Increased persistence index indicates less fatigability to the step illumination. Persistence index increased starting 2 d post-lesion in 7/9 mice to 254% of baseline values (**Fig. 3C, right**) (0.3012 to 0.7655; ANOVA of persistence index over days, F=27.73, p<2.2e-16). Results for 10 Hz pulsed illumination were similar to the 1000 ms step illumination (**Supplemental Figure 1**; goo.gl/sWRNLU). Further comparison of 7 d and pre-lesion response duration families showed increased peak amplitudes especially for short duration stimuli (**Fig. 3D, left**), indicating reduced threshold due to increased sensitivity. Similar or even more dramatic effects were evident for peak velocity (**Fig. 3D, middle**). Parallel experiments in the same mice tracked whisker retractions evoked by illumination of the caudal retraction area (Park et al 2016) and found increased sensitivity, but not amplitude or velocity, and minimal changes in fatigability (**Supplemental Fig. 2**; goo.gl/3SDp8q), suggesting that many of the results are unique to whisker protractions. Finally, we summarized the effects of denervation on nerve- and muscle-evoked whisker protractions by comparing the changes in peak amplitude and peak velocity (24 h in ChAT-ChR2 and 7 d in Emx1-ChR2 mice). Nerve- and muscle-evoked whisker movements change in opposite directions after facial nerve transection: nerve-evoked responses were abolished by 24 h, while muscle-evoked responses increased in amplitude and velocity (**Fig. 3D, right**). The velocity of pre-lesion optogenetic muscle-evoked whisker movements was less than nerve-evoked movements (t-test of the peak velocity of pre-lesion ChAT-ChR2 and pre-lesion Emx1-ChR2 mice, t=3.21, p=0.007), but increased at 7 d post-lesion to approximately 1000°/s, similar to the fastest pre-lesion nerve-evoked whisker movements (t-test of the peak velocity of pre-lesion ChAT-ChR2 and 7 d Emx1-ChR2 mice, t=0.28, p=0.78). Together these results indicate that denervated whisker pad protraction muscles undergo marked increases in movement capacity, including increased amplitude, velocity and sensitivity, and decreased fatigability.

**Figure 3.**
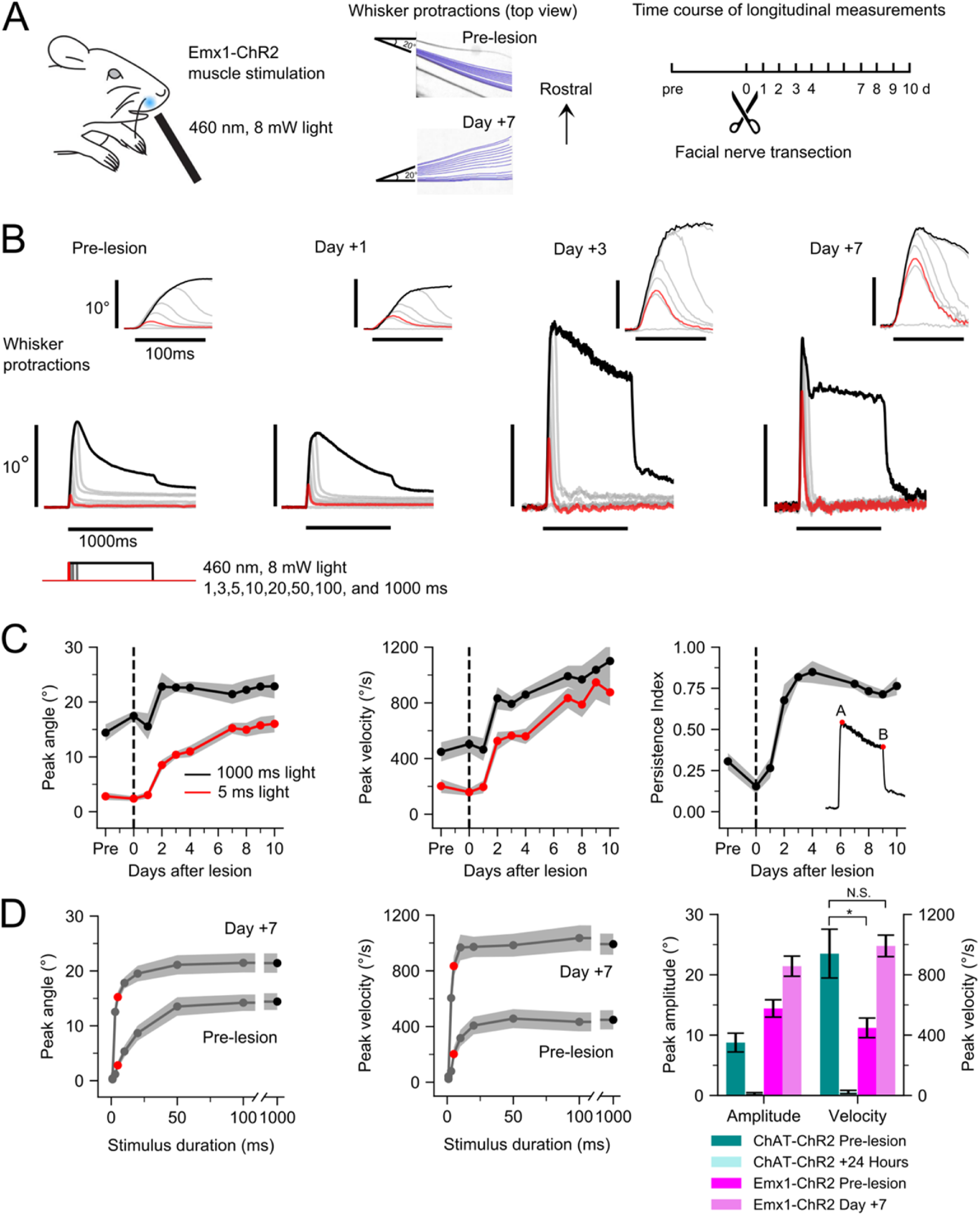
Functional enhancement of denervated muscle. **(A)** Optogenetic muscle paradigm. Left: Schematic of transdermal muscle illumination in Emx1-ChR2 tion of optogenetic illumination indicated with blue spot. Middle: Frame-by-frame position of one tracked whisker in an example subject in response to a 1000 ms, 8 mW pulse of 460 nm LED illumination pre-lesion and 7 d post-lesion. Right: Experimental timeline. **(B)** Whisker protractions evoked by optogenetic muscle stimulation from an example mouse at various times relative to nerve transection. Each trace is the average of 10 responses to a different duration of illumination at 8 mW. Stimuli are 1, 3, 5, 10, 20, 50, 100, 1000 ms, with 5 ms (red) and 1000 ms (black) highlighted. Insets show the same traces on an expanded time scale. **(C)** Time course of peak amplitude (left) and peak velocity (middle) of whisker protractions in responses to 5 ms (red) and 1000 ms (black) light stimuli. Time course of persistence index for 1000 ms stimulus (right); inset shows that persistence index is the ratio of response amplitude at 1000 ms to peak amplitude (B/A). Values are mean ± SEM; n=9 mice. **(D)** Pre-lesion and 7 d response duration families for peak amplitude (left) and peak velocity (middle). Red circle, 5 ms; black circle, 1000 ms stimuli. Right: Summary data comparing amplitude and velocity changes for ChAT-ChR2 (blues) and Emx1-ChR2 mice (magentas) in response to 1000 ms stimuli. T- tests compare velocity values for pre-lesion ChAT-ChR2 with pre-lesion Emx1-ChR2, and pre-lesion ChAT-ChR2 with day + 7 Emx1-ChR2. Values are mean ± SEM; n=9 mice. * indicates p<0.05, N.S. indicates not significant. ranscriptome of denervated whisker pad muscle.

### Transcriptome of denervated muscle

To determine transcriptional changes underlying denervation-induced enhanced muscle function, we performed RNA-seq analysis of denervated and sham-denervated whisker pad tissue at 1, 3, and 7 d after facial nerve lesion. We calculated the differential expression of transcripts using the Kallisto/Sleuth pipeline (Bray et al 2016) and gene ontology (GO) using clusterProfiler (Pimentel et al 2017). Differentially expressed transcripts were evaluated with an experiment-wide Likelihood Ratio Test (LRT) to account for multiple measurements. Injury- and day-specific Wald test results that showed at least a 2-fold change (FC), and were filtered by transcripts previously identified as differentially expressed from LRT, were used to classify the changes in transcript levels across days (Methods). After denervation, 389 transcripts were differentially expressed in whisker pad, with increased expression between 1 to 3 and 7 d (**Fig. 4A**). We compared these whisker pad data with a published RNA-seq dataset from the atrophy-prone soleus muscle at 3 d after denervation (Macpherson et al 2015). 759 transcripts were determined to be differentially expressed in injured soleus. 63 of 692 transcripts were commonly regulated between whisker pad and soleus at 3 d after denervation, 629/692 were unique to one of the two models, and no transcripts were regulated with at least a 2 FC in opposite directions (**Fig. 4B**). Notably, there was limited overlap in GO terms and many differences (**Fig. 4C**). We further analyzed genes from the cell component ontology, focusing on selected terms of interest. These are shown as heatmaps of normalized expression levels at days 1, 3 and 7 d for whisker pad and day 3 for soleus (transcript levels were normalized by z-score across all conditions). While genes related to atrophy and cholinergic receptors, for example, were similarly upregulated in whisker pad and soleus muscle, there were a number of differences, such as the expression of key ion channel and contractile fiber genes (**Fig. 4D**). More complete lists are presented in **Supplemental Fig. 3** (goo.gl/Afe8BE) and the full dataset can be found at GEO with accession GSE121590. These comprehensive transcriptome-level data from denervated whisker pad muscle represent a key basis for further understanding the enhanced functional changes we have observed.

**Figure 4.**
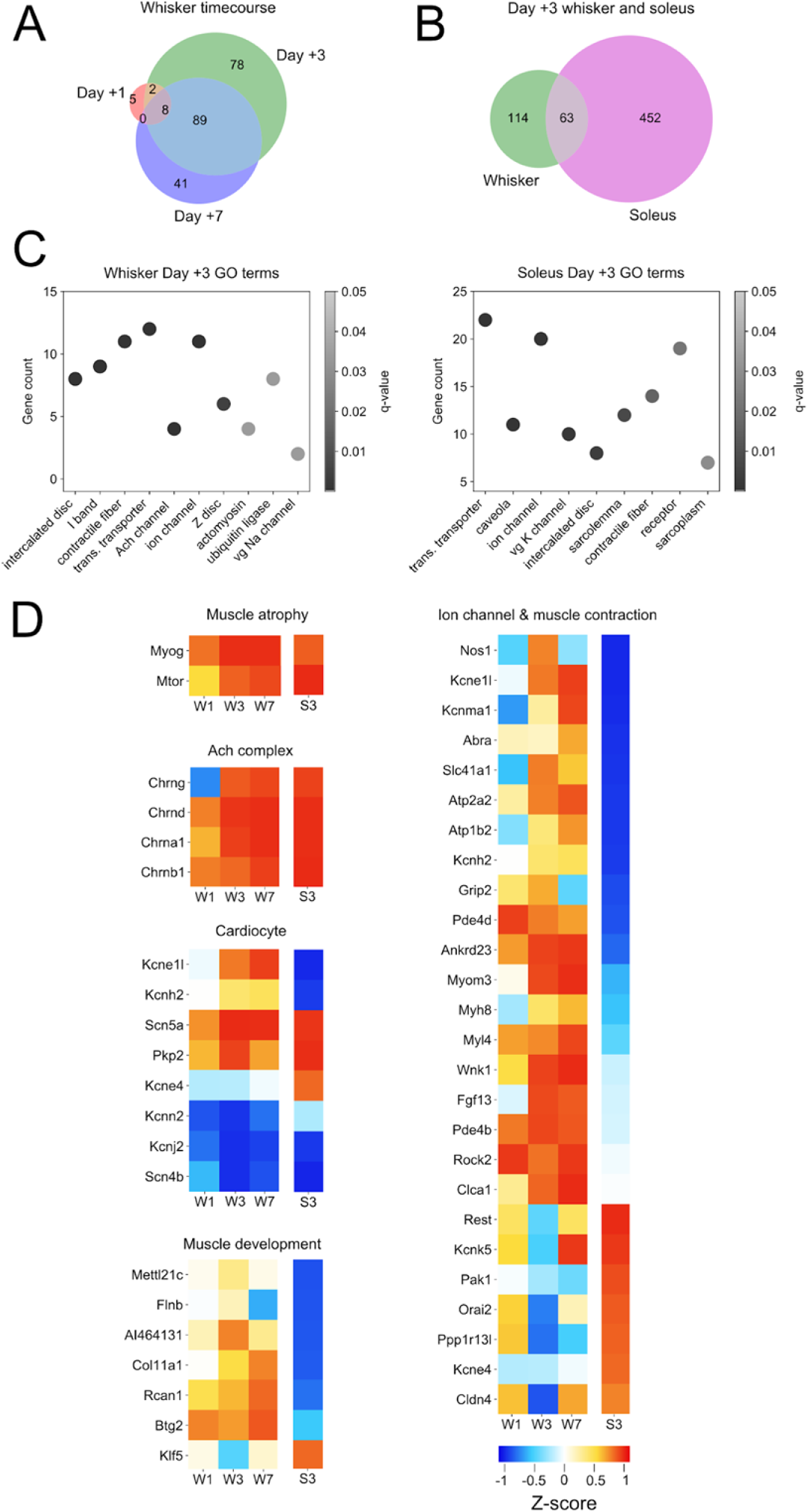
Transcriptome of denervated whisker pad muscle. **(A)** Venn diagram of differentially expressed transcripts from whisker pad with at least ± 2-fold change (FC) at 1, 3 and 7 d post-denervation (n=3 mice at each time point). **(B)** Venn diagram of differentially expressed transcripts with ± 2-FC from whisker pad and soleus at 3 d post-denervation. Soleus data in panels B, C and D is from (Macpherson et al., 2011; n=2 mice). **(C)** Subset of enriched cell component GO terms for whisker pad (left) and soleus (right) at 3 d post-denervation. Identity numbers of GO terms are listed in Methods. **(D)** Heatmaps of expression for gene families related to Muscle atrophy, ACh complex, Cardiocyte, Muscle development, and Ion channels & Muscle contraction. Data shown as average z-score. Columns labeled W1, W3 and W7 indicate whisker pad data from 1, 3, and 7 d post-denervation, respectively. Rightmost columns labeled S3 indicate soleus data from 3 d post denervation. For Muscle development and Ion channel & Muscle contraction groups, genes included in heatmap were limited to those that had polar directional z-scores between whisker pad and soleus at 3 d.

## DISCUSSION

We determined the functional and transcriptomic changes of mouse whisker pad that result from facial motor nerve denervation. To our knowledge, our results represent the first demonstration of enhanced functional properties of paralyzed whisker pad muscles, and provide a novel RNA-seq dataset of the transcriptomic changes underlying this process. Our study has important implications for developing future treatments for the paralysis that occurs in debilitating disorders such as nerve damage and lower motor neuron degenerative diseases.

### Functional changes in nerve and muscle revealed by peripheral optogenetic stimulation

In the days following motor nerve transection, the distal cut nerve undergoes a process of Wallerian degeneration, leaving neuromuscular synaptic terminals transiently intact before complete denervation (Tricaud & Park 2017). We used the ability to non-invasively induce whisker movements longitudinally over days to measure changes in the functional properties of facial nerve and whisker pad muscle throughout this process. Optogenetic stimulation of the distal cut end of the facial nerve continued to produce movements for up to 24 h but degraded thereafter, consistent with the time course of Wallerian degeneration (Tricaud & Park 2017). After 24 h, the functional changes that occur in muscle can be attributed to the intrinsic muscle changes in response to the denervation-induced loss of cholinergic input. Most muscles respond to this situation with atrophy and loss of function, as evidenced by a large body of work (Carlson 2014, Thesleff 1974, Wu et al 2014). A recent study that used optogenetic stimulation of triceps surae muscle found more than 50% loss of both force generation and cross-sectional area by 3 d after sciatic nerve lesion (Magown et al 2015). Based on these results, we expected that facial nerve lesion would result in loss of whisker pad muscle function and reduced whisker movements. However, we found the opposite effect. Two days after facial nerve transection, the same optogenetic light stimuli applied to whisker pad muscle produced large amplitude and faster whisker movements with less fatigue. Remarkably, brief light stimuli that produced no movements before denervation became highly effective. For example, the response to a 5 ms stimulus became near maximal and indistinguishable from the strongest (1000 ms) stimulus, whereas there was no response to the 5 ms stimulus before denervation. These gain-of-function changes lasted at least 10 d after denervation (the length of our study) with no indication of reversal, suggesting that it is unlikely we simply missed a delayed loss-of-function response. Rather, it is more likely that whisker pad muscles respond to denervation in a distinct way compared to other muscle types. Our gene profiling results (below) reveal many differences with the atrophy-prone soleus muscle, further supporting this idea. Importantly, expression levels of ChR2 from whisker pad tissue in Emx1-ChR2 mice did not increase after facial nerve transection (Supplemental Fig. 4; goo.gl/sb75Qd), ruling out the possibility that increased ChR2 expression led to the functional changes that we observed.

### Fasciculations, atrophy, and excitability

Fasciculations (also known as fibrillations) are small-amplitude spontaneous muscle twitches that commonly occur after denervation (Wu et al 2014). Fasciculations are caused by oscillations in the membrane potential of muscle cells that result from denervation-induced changes in the intrinsic excitability of muscle cells, including tonic membrane depolarization and altered sodium and potassium conductances (Salafsky et al 1968, Thesleff & Ward 1975). In the rodent whisker system, fasciculations can be precisely measured after facial nerve transection by noninvasive video monitoring of the paralyzed whiskers, and furthermore are a useful indicator of the state of denervation because they are abolished by reinnervation (Heaton & Kobler 2005). It is tempting to speculate that fasciculations could help protect denervated muscle from atrophy by providing constant excitation, but because fasciculations occur even in muscles that show loss of function, the relationship between fasciculations and resilience is still unclear and may vary between different muscle types. It is possible that the strength of fasciculations is greater in whisker pad compared to other muscles, or that additional functional or genetic factors are involved in counteracting loss of function. The relationship between fasciculations, excitability, and resilience to atrophy will need to be determined in future studies.

### Strategies for stimulation of paralyzed muscle

Only a few studies have demonstrated direct control of muscle via muscle-specific expression of optogenetic actuators (Bruegmann et al 2015, Bryson et al 2014, Magown et al 2015, Park et al 2016). The optogenetic approach has major advantages over currently available technology for restoration of movement. One of the advantages is the direct and safe stimulation of muscle in the absence of nerve. This is different than functional electrical stimulation (FES), a commonly used method in the clinic (Doucet et al 2012, Ho et al 2014, Peckham & Knutson 2005), because FES depends on nerve rather than muscle stimulation. A critical limitation of FES is the presence and viability of nerve, which may be compromised in cases of severe damage or degenerative diseases. The high electrical currents needed in these cases can cause problems such as tissue damage, non-specific spread across muscle groups, and discomfort (Plenk 2011). Thus, the ability of optogenetics to non-invasively stimulate muscle in the absence of nerve is a major advantage compared to current electrical approaches or optogenetic nerve stimulation approaches (Llewellyn et al 2010), and indeed could be the only viable approach to restore the function of muscle after severe injuries or diseases that result in complete and permanent denervation. There are still significant issues that need to be solved in order to translate optogenetics to the clinic, including the expression of light-sensitive proteins in tissue of interest using viral vectors and appropriate hardware for light delivery (Fenno et al 2011, Montgomery et al 2016, Yizhar et al 2011), and thus these remain active areas of research.

### Transcriptome of denervated muscle

Discovering the expression changes that occur in denervated muscle is an important goal for developing strategies to restore muscle function. We used RNA-seq to detect and quantify transcript levels changing over time following denervation. In particular, our objective was to understand the unique transcriptomic landscape of the denervated whisker pad, which shows enhanced muscle function, compared to other muscle samples that show loss of function. Thus, by comparing our dataset with an existing dataset from atrophy-prone soleus muscle at a matching 3 d time point after denervation (Macpherson et al 2015), we identified unique features of the denervated whisker pad. Our results indicate that expression of several key genes was regulated in the same direction in both whisker pad and soleus (either up- or both down-regulated), while many others showed differential expression as we would have predicted based on functional differences. For example, expression of the atrophy-related genes *Myog* and *Mtor*, and *Chrn* nicotinic cholinergic receptor family genes were upregulated in both whisker pad and soleus. However, there were prominent differences in expression of genes for ion channels and contractile fibers. For example, the *Kcn* family of voltage-gated potassium channels showed several instances of upregulation in whisker pad and downregulation in soleus, and vice versa. Similarly, slow-twitch (type II) contractile fibers, including *Myom3* and *Myh8*, were upregulated in whisker pad and downregulated in soleus, consistent with fiber-type switches after denervation. It is likely that the differences in expression of ion channels and contractile fibers at least partly explain the distinct functional responses of these muscles to denervation. For example, the upregulation of slow-twitch fibers in whisker pad is consistent with the striking decrease in fatigability that we observed (Figure 3). Changes in expression of voltage-gated ion channels, including potassium channels, sodium channels, and many others, can result in membrane potential depolarization, oscillations, and changes in the waveform of the action potential. These features likely relate to the changes in optogenetically evoked whisker movements that we measured, including increased sensitivity, amplitude, and velocity. The detailed relationship between ion channel and contractile fiber expression with the functional changes requires further investigation. Another outstanding question is how changes in excitability relate to susceptibility to atrophy. One possibility is that atrophy-related genetic programs are activated in all denervated muscle types, but additional changes in excitability that lead to enhanced function in some muscles types essentially counteract this process. Since our experiments focused on functional changes in muscle responses as measured with optogenetic stimulation and video recording of whisker movements, further studies are needed to determine whether atrophy occurred on the level of individual muscle fibers.

It should be noted that our samples were limited to the rostral protraction area of the whisker pad. At this location, the spot of light activates primarily the extrinsic protractor muscles pars media superior and pars media inferior of *M. nasolabialis profundus* and also the intrinsic follicular muscles (Haidarliu et al 2010, Jin et al 2004, Park et al 2016). In the future, it would be interesting to compare RNA-seq results with the caudal retraction area, which showed increased sensitivity but more subtle changes in amplitude and fatigability (Supplemental Figure 2; goo.gl/3SDp8q), to identify further links between changes in gene expression and movement phenotypes. It would also be important in future work to obtain more complete datasets comparing functional and transcriptomic results from several muscles types. This would help address drawbacks in the comparison of our RNA-seq dataset with published data from soleus (Macpherson et al 2015), including that the samples were obtained from mice of different ages, and that our biopsied samples included both skin and muscle tissue. Future work could also directly relate transcriptomic and functional data by performing genetic analysis of the same subjects used for the optogenetic measures, which was not done here.

## Conclusion

In conclusion, our combined functional and transcriptomic results have implications for future work on the restoration of movement following nerve damage or motoneuron degeneration. The unique properties of denervated whisker pad muscles that we have identified could inform future genetic strategies to restore muscle function after nerve injury or motoneuron degeneration in various neuromuscular systems.

## Acknowledgements

Supported in part by grants from the National Institutes of Health (R01NS094450) and the New Jersey Commission on Brain Injury Research (CBIR16IRG032) to DJM. AU was a member of the NeuroSURP program (NIH R25NS105143). We thank S. Olga Yiantsos, Ridhima Sakhuja, Fizza Fahim, Sydney Terris, Hill Chang, Jennifer Fang and Emefa Ocansey for assistance with experiments. We thank RUCDR Infinite Biologics staff for collection of RNA-seq data and the Office of Advanced Research Computing (OARC) at Rutgers, The State University of New Jersey for providing access to the Perceval cluster and associated research computing resources that have contributed to the results reported here with support from NIH (S10OD012346). URL: http://oarc.rutgers.edu. Author contributions: AB, TJV, AU, RPH, CRL, DJM designed experiments, AB, AU, CRL, MRS, DJM performed experiments, TJV, AB, RPH analyzed data, DJM drafted manuscript with input from all authors.

---

Data supplements are available via links in text and https://github.com/margolislab/Vajtay_Bandi_2018.

**Supplemental Figure 1.**
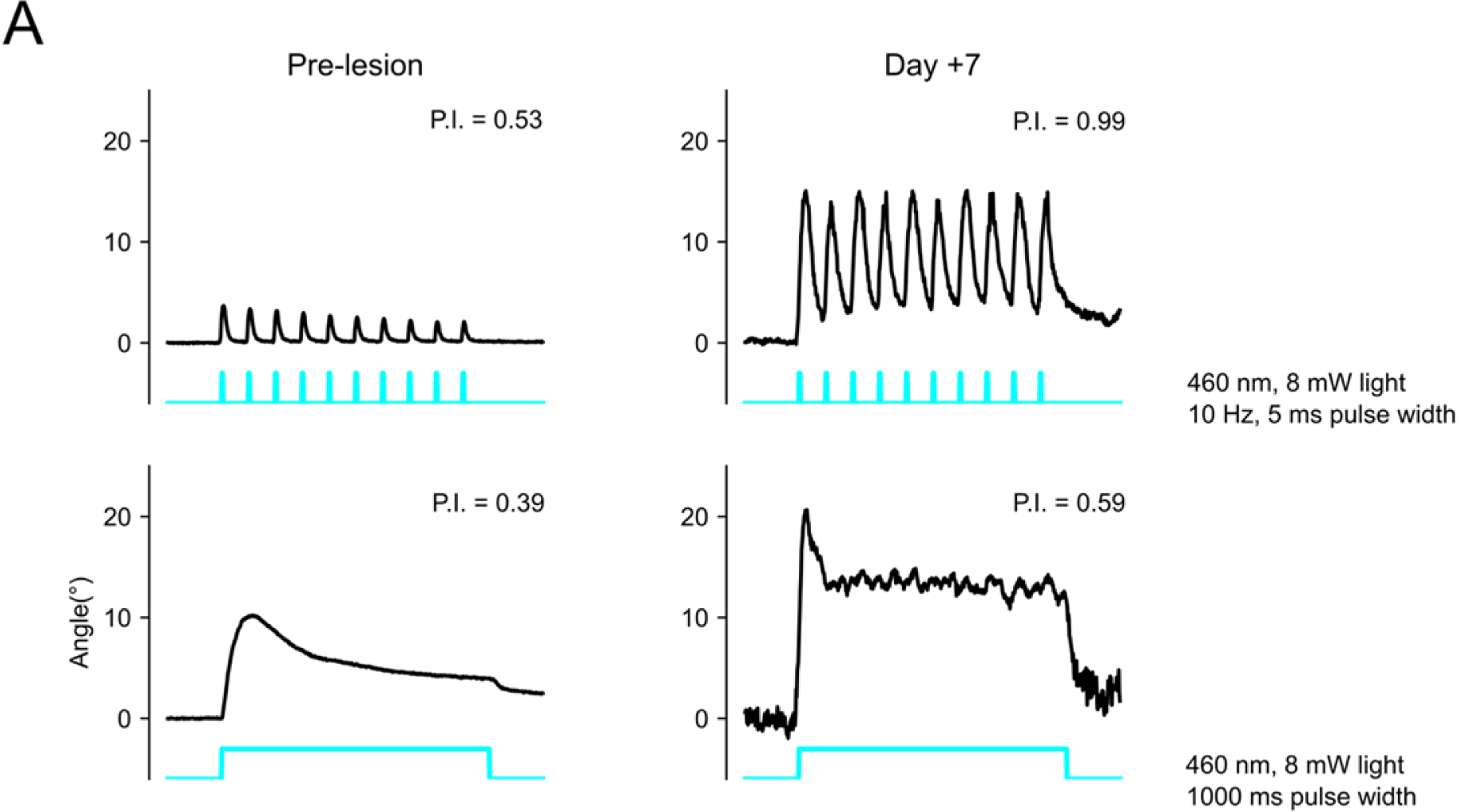
Comparison of whisker protractions to 10 Hz and step illumination. **(A)** Top row shows responses to 10 Hz light stimuli (5 ms pulse width) before and 7 days after nerve transection in an example mouse. Bottom row shows responses to step illumination (1000 ms) before and 7 days after nerve transection in the same example mouse as in the top row. Each trace is a representative single trial.

**Supplemental Figure 2.**
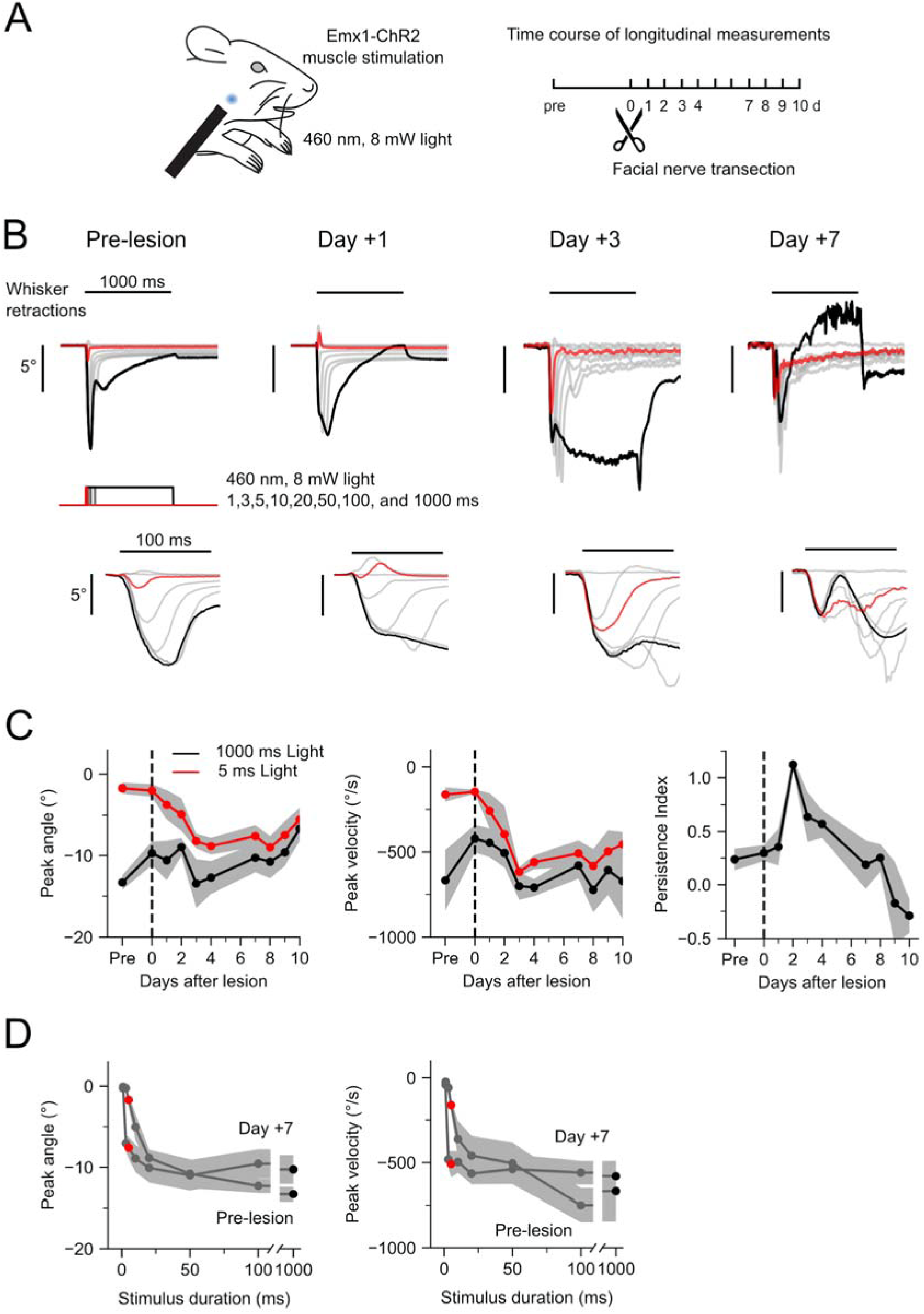
Effects of denervation at caudal retraction area. **A)** Optogenetic muscle paradigm. Left: Schematic of transdermal muscle illumination in Emx1-ChR2 tion of optogenetic illumination indicated with blue spot. Right: Experimental timeline. **(B)** Whisker retractions evoked by optogenetic nerve stimulation from an example mouse at various times relative to nerve transection. Each trace is the average of 10 responses to a different duration of illumination at 8 mW. red, 5 ms stimulus; black, 1000 ms stimulus; gray, 1, 3, 10, 20, 50, 100 ms. Insets (below) show movements on an expanded time scale. **(C)** Time course of peak amplitude (left), time course of peak velocity (middle), and time course of persistence index from 1000 ms stimuli. High persistence index indicates low fatigability. Red, 5 ms; black, 1000 ms stimuli. Values are mean ± SEM; n=9 mice. **(D)** Comparison of baseline and 7 d response duration families for peak amplitude (left) and peak velocity (middle). Red, 5 ms; black, 1000 ms stimuli. Values are mean ± SEM; n=9 mice

**Supplemental Figure 3.**
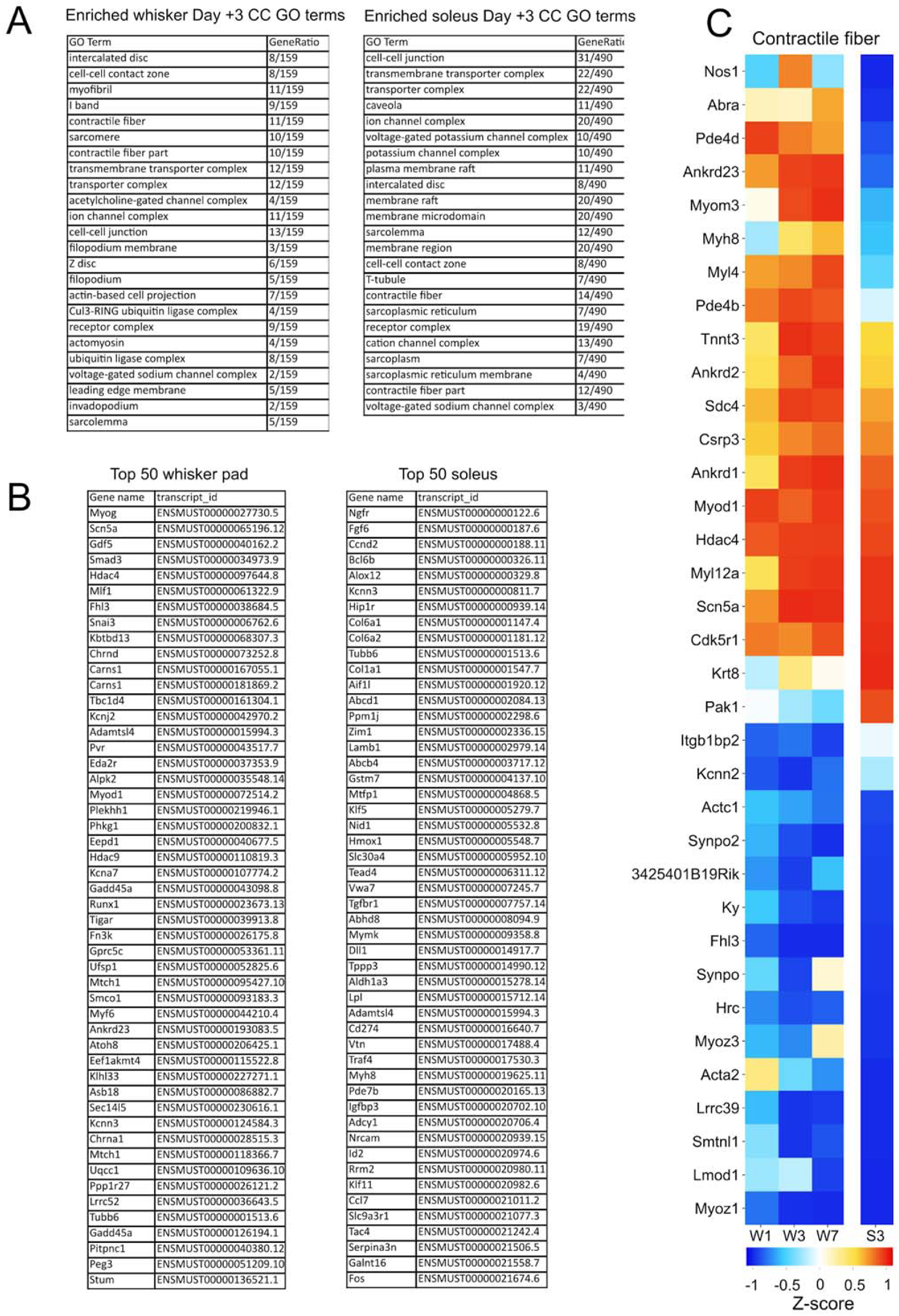
Additional transcriptome data for denervated whisker pad and soleus muscle. **(A)** Full lists of enriched GO terms for both whisker pad and soleus 3 d calculated from a ranked list of differentially expressed ± 2-FC transcript gene IDs. All GO terms listed were significantly enriched by hypergeometric test at q<0.05. **(B)** List of top 50 significant differentially expressed transcripts for both whisker pad and soleus. **(C)** Full heatmap of differentially expressed genes included in the Contractile fiber GO term. Data shown as average z-score. Columns labeled W1, W3 and W7 indicate whisker pad data from 1, 3, and 7 d post-denervation, respectively. Rightmost columns labeled S3 indicate soleus data from 3 d post denervation.

**Supplemental Figure 4.**
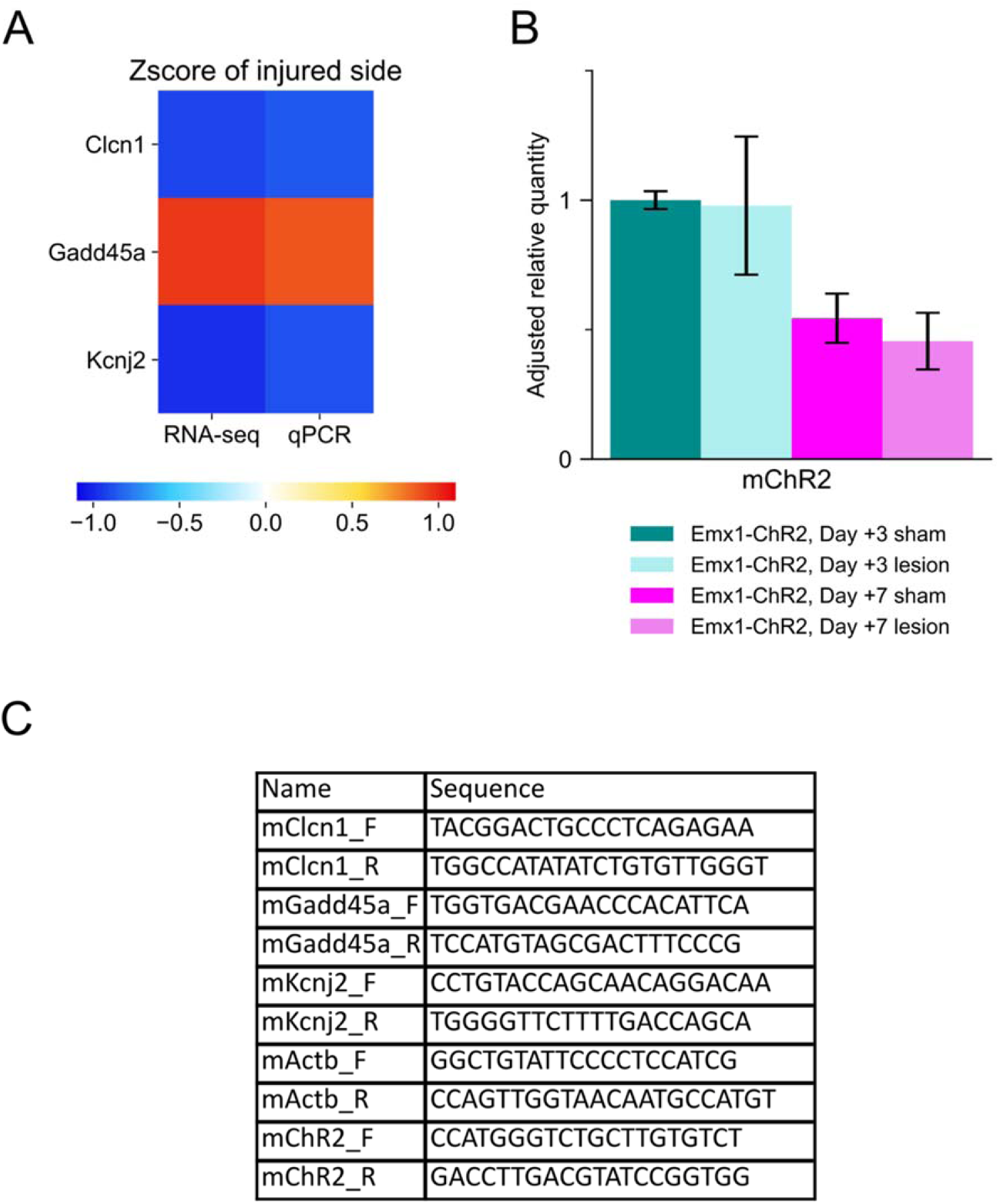
qPCR validation of RNA-seq data and measurements of ChR2 expression. **(A)** Heatmap of differential expression of Clcn1, Gadd45a, and Kcnj2 from 3 d whisker pad for RNA-seq (left column) and qPCR (right column). qPCR values were normalized to ActB expression. **(B)** qPCR data comparing ChR2 expression levels in sham and lesioned whisker pad tissue from Emx1-ChR2 mice. Bars (errorbars) represent mean (± SEM) data from left (sham) and right (lesion) whisker pads from day 3 (n=3 mice) and day 7 (n=2 mice) after lesion. One of the five subjects included was unlesioned. No increases were detected at 3 d and 7 d post-lesion. **(C)** Name and sequence of primers used for qPCR.

